# Selective disruption of Traf1/cIAP2 interaction attenuates inflammatory responses and limits sepsis and rheumatoid arthritis

**DOI:** 10.1101/2024.08.31.610651

**Authors:** Yitian Tang, Fatemah Aleithan, Sahib Singh Madahar, Ali Mirzaesmaeili, Sunpreet Saran, Jialing Tang, Safoura Zangiabadi, Robert Inman, Gary Sweeney, Ali A. Abdul-Sater

## Abstract

Tumor necrosis factor receptor-associated factor 1 (TRAF1) is an immune signaling adapter protein linked to increased susceptibility to rheumatoid arthritis (RA). TRAF1 has dual roles in regulating NF-κB and MAPK signaling: it promotes signaling through its association with cellular inhibitor of apoptosis 2 (cIAP2) downstream of certain tumor necrosis factor receptor (TNFR) family members but inhibits Toll-like receptor (TLR) signaling by limiting linear ubiquitination of key signaling proteins. Here, we identify a critical mutation in TRAF1 (V203A in humans, V196A in mice) that disrupts its interaction with cIAP2, leading to a significant reduction in TLR signaling and downstream inflammation in human and murine macrophages. We demonstrate that TRAF1 is recruited to the TLR4 complex and is indispensable for the recruitment of cIAP2, facilitating TAK1 phosphorylation and the activation of NF-κB and MAPK signaling pathways. Remarkably, mice harboring the TRAF1 V196A mutation are protected from LPS-induced septic shock and exhibit markedly reduced joint inflammation and disease severity in a collagen antibody-induced arthritis (CAIA) model of RA. These findings reveal a previously unrecognized and crucial role for the TRAF1/cIAP2 axis in promoting inflammation and offer a promising foundation for the development of novel therapeutic strategies for inflammatory conditions, such as sepsis and RA.

## INTRODUCTION

Rheumatoid arthritis (RA), though multifactorial, is characterized by misdirected and overactive immune/inflammatory responses leading to joint damage. The aberrant immune responses in RA involve the activation of various immune cells and inflammatory molecules (*1*, *2*). T lymphocytes and macrophages play a significant role in perpetuating and driving the disease. When activated, these cells produce mediators, including cytokines, that activate resident cells such as synovial fibroblasts, osteoblasts, and osteoclasts. These resident cells, in turn, secrete more pro-inflammatory cytokines and chemokines, which recruit additional immune cells into the inflamed joint, exacerbating the inflammatory response (*1*). This understanding has led to the development of new drugs, including biologics and small-molecule inhibitors, that block these cytokines (e.g., TNF blockers) and associated signaling pathways (*3*). Unfortunately, these therapies are only partially effective or completely ineffective in nearly half of the patients, likely due to the involvement of multiple cytokines and signaling pathways in disease pathogenesis. There is an urgent need for therapies that specifically target the immune and inflammatory components driving RA pathogenesis and “pump the brakes” on their ability to produce damaging cytokines.

In addition to signaling through their antigen receptors, lymphocytes require signals from Tumor Necrosis Factor Receptor superfamily (TNFR) members such as 4-1BB and CD40 in T and B cells, respectively. These receptors activate NF-κB and MAPKs to induce survival, proliferation, and cytokine production (*4*). The adapter protein TNFR-associated factor 1 (TRAF1), along with TRAF2, recruits cellular inhibitor of apoptosis protein 2 (cIAP2) to activate NF-κB and other MAPK signaling pathways (*5–12*). Importantly, several TNFRs have been implicated in RA pathogenesis, and therapies targeting them are being considered (*13*). We recently demonstrated that reducing TRAF1 levels, which are overexpressed in 48% of B cell-related cancers, decreases the activation of NF-κB and other survival pathways, leading to the death of B leukemia cells from chronic lymphocytic leukemia (CLL) patients (*14*). Conversely, in another study, we showed that in addition to activating B and T cells, TRAF1 plays an opposing role in monocytes and macrophages by limiting NF-κB activation and reducing cytokine production and inflammation (*15*). We found that in macrophages, which rely on Toll-like receptors (TLRs) to induce pro-inflammatory cytokines, TRAF1 directly interacts with and disrupts the linear ubiquitin chain assembly complex (LUBAC), thereby limiting linear ubiquitination of IKKγ (NEMO) and reducing NF-κB activation (*15*). This explains findings from genome-wide association studies (GWAS) that identified single nucleotide polymorphisms (SNPs) in TRAF1 as being associated with an increased risk of developing RA (*16–19*) and juvenile idiopathic arthritis (*20–22*). Additionally, RA patients with this SNP are at a higher risk of mortality from sepsis (*19*). We found that individuals with the TRAF1 variant that increases susceptibility to RA have lower TRAF1 levels in their T lymphocytes and macrophages. While their T lymphocytes were less activated and produced lower cytokine levels, their monocytes produced markedly higher cytokine levels, ultimately leading to a hyper-inflammatory state that may drive RA pathogenesis. We also showed that TRAF1-deficient mice are highly susceptible to LPS-induced sepsis (*15*).

In a more recent study, we demonstrated that intraarticular injection of TRAF1-deficient macrophages into the knee joints of wildtype (WT) mice subjected to the collagen antibody-induced arthritis (CAIA) model of RA led to exacerbated joint inflammation and cellular infiltration compared to WT mice injected with WT macrophages (*23*). Moreover, we showed in another study that TRAF1 can limit the secretion of the potent pro-inflammatory cytokine IL-1β independently of NF-κB inhibition by reducing the assembly of the inflammasome complex and attenuating the linear ubiquitination of the adapter protein ASC (*24*). We also showed that TRAF1 knockout mice developed more severe gout, a common form of inflammatory arthritis primarily driven by inflammasome activation (*24*). Therefore, TRAF1 can have opposing roles in regulating NF-κB activation based on the cell type and receptor signaling pathway. Here, we show that by mutating a single amino acid in TRAF1 (V203A), we disrupt its association with cIAP2, resulting in significant attenuation of LPS-induced inflammation in macrophages. Remarkably, we generated mice harboring this mutation (TRAF1 V196A) and showed that these mice are protected from LPS-induced sepsis and exhibit markedly diminished joint inflammation and disease severity in the CAIA model of RA.

## RESULTS

### V203 is required for TRAF1’s interaction with cIAP2

To dissect the dichotomous role of TRAF1 in NF-κB signaling and specifically inhibit the TRAF1/cIAP2/NF-κB axis without disrupting TRAF1-mediated attenuation of linear ubiquitination, we employed site-directed mutagenesis to identify amino acid residues critical for the interaction between TRAF1 and cIAP2. Strikingly, we found that a single mutation of either valine at position 203 (V203) or glutamate at position 207 (E207) to alanine (V203A or E207A) in the TRAF1 protein largely abrogates its interaction with cIAP2 (**Fig. 1a**). Importantly, these mutations did not affect TRAF1’s interaction with HOIL-1, a component of LUBAC, and nearby mutations (e.g., E209A) did not affect the TRAF1/cIAP2 interaction (**Fig. 1b**). Crucially, the V203A and E207A mutations did not alter TRAF1’s interaction with another key signaling partner, TRAF2 (**Fig. 1c**).

**Figure 1:**
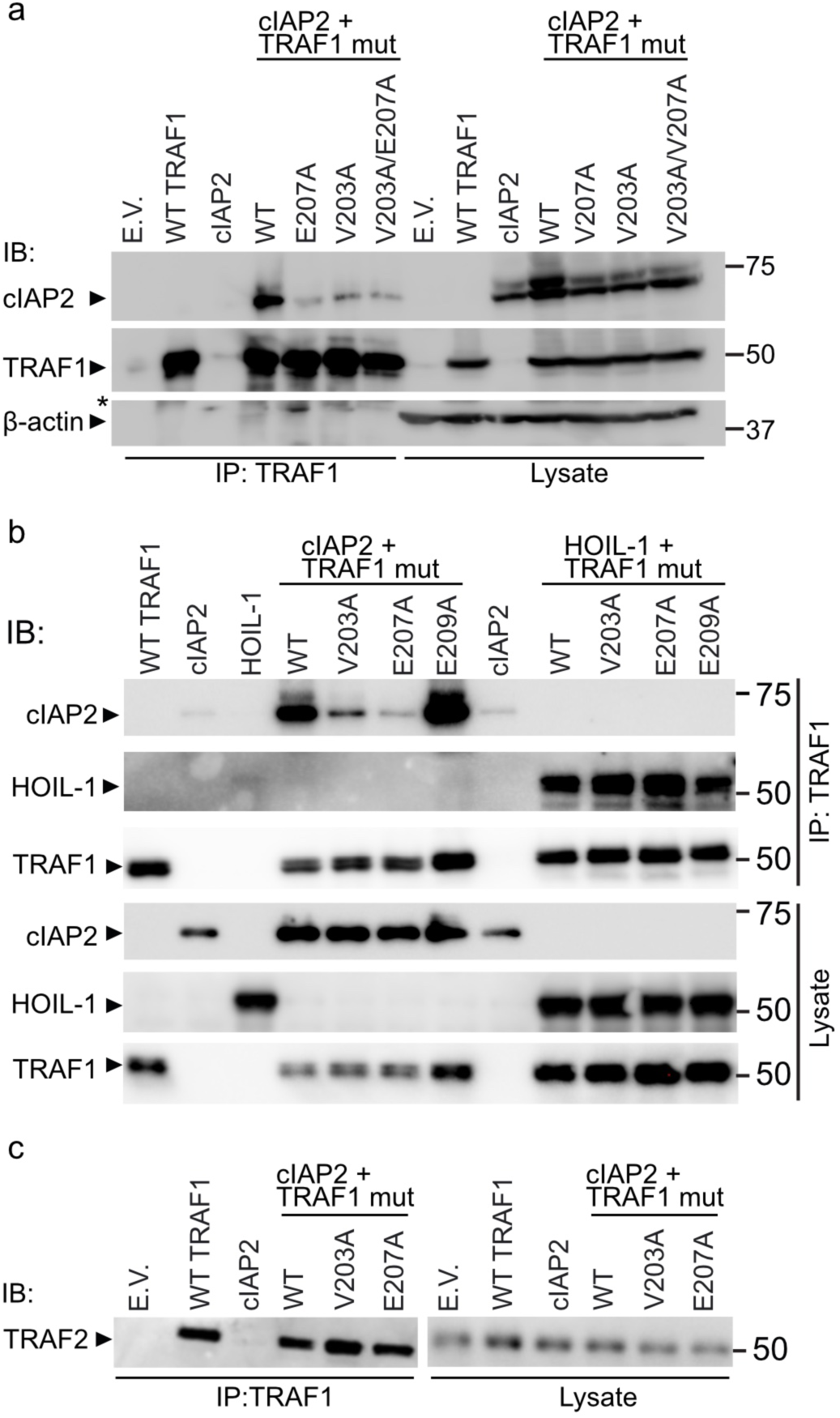
TRAF1 V203 and V207 are required for interaction with cIAP2. (**a**) Immunoprecipitation (IP) of TRAF1 from lysates of HEK 293T cells overexpressing cIAP2 with or without wildtype (WT) or various TRAF1 mutant constructs as indicated and then blotted for cIAP2, TRAF1, or as a loading control β-Actin. (**b**) Immunoprecipitation (IP) of TRAF1 from lysates of HEK 293T cells overexpressing cIAP2 or HOIL-1 with or without various TRAF1 mutant constructs as indicated and then blotted for cIAP2, HOIL-1 or TRAF1. (**c**) TRAF1 IP of HEK 293T lysates, as in panel a, were probed with blotted for TRAF2. E.V. empty vector. All blots are representative of three independent experiments.

### TRAF1 protects cIAP2 from degradation

Intriguingly, we observed that when TRAF1 is overexpressed with cIAP2, cIAP2 levels increased in 293T cells where TRAF1 signaling does not induce NF-κB activation. To investigate this, we performed a cycloheximide chase assay and demonstrated that cIAP2 stability was increased when co-expressed with TRAF1 (**Fig. 2a**). Consistently, in RAJI lymphoma cells that naturally express high levels of TRAF1, endogenous cIAP2 stability was lower when TRAF1 was knocked down (**Fig. 2b**). Importantly, the interaction with TRAF1 was required for increasing cIAP2 stability because we show that overexpression of WT TRAF1, but not V203A or E207A TRAF1 mutants, prolongs the half-life of cIAP2 (**Fig. 2c, d; supplementary Fig. 1**). Mechanistically, cIAP2 degradation was mediated by the proteasome (**Fig. 2e**). Since cIAP2 is known to be auto-ubiquitinated and cross-ubiquitinated by cIAP1, we overexpressed ligase-dead cIAP2 (H574A) and cIAP1 (H588A) and demonstrated that TRAF1 overexpression increased cIAP2 stability only when cIAP1, but not cIAP1 H588A, was co-expressed (**Fig. 2f**). This indicates that TRAF1 protects cIAP2 from cIAP1-mediated degradation. Collectively, these results identify V203A and E207A mutations as strong candidates to specifically interfere with TRAF1/cIAP2 interaction and downstream signaling without affecting its association with LUBAC or other proteins.

**Figure 2:**
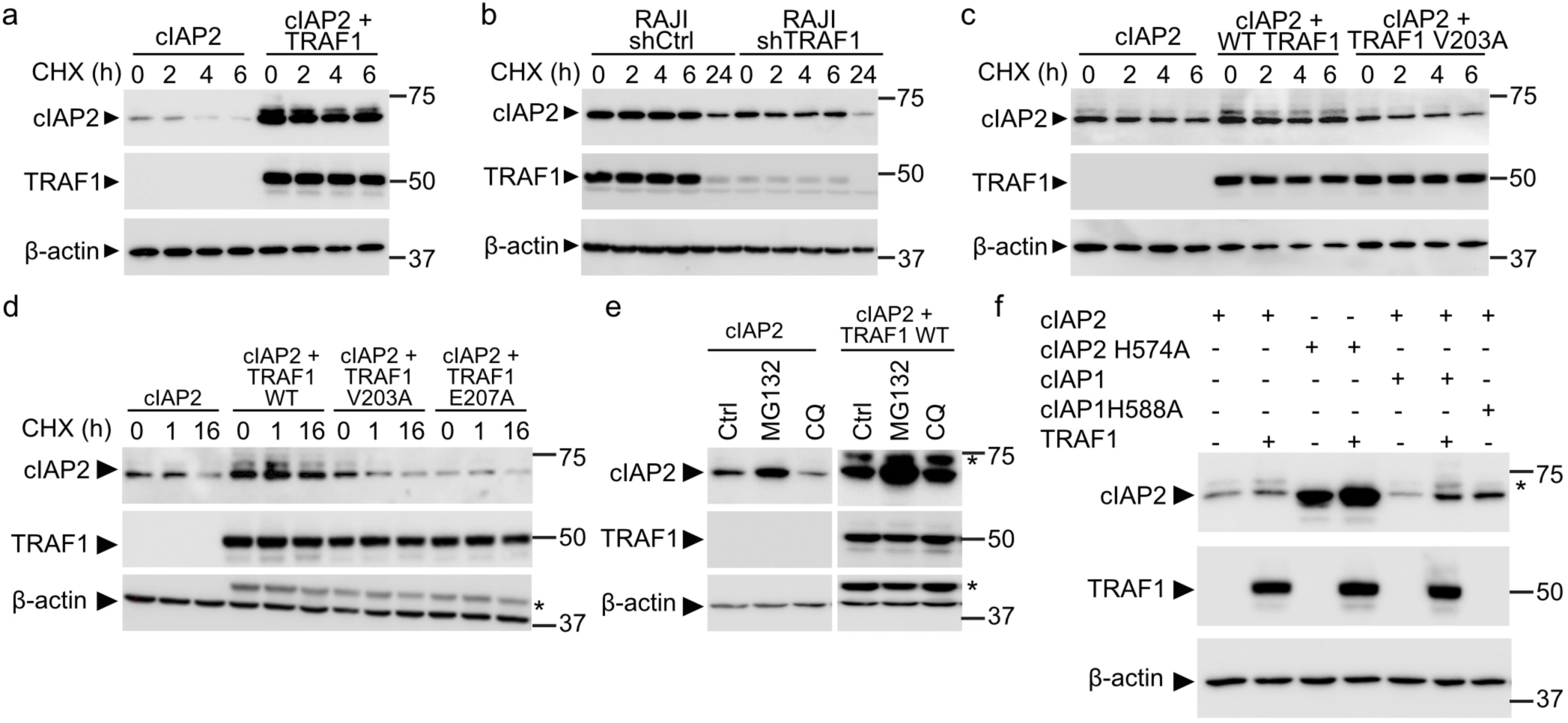
TRAF1 stabilizes cIAP2 and protects it from degradation. (**a**) HEK 293T cells overexpressing cIAP2 with or without WT TRAF1 were treated with 5 µg/ml cycloheximide (CHX) to inhibit denovo protein synthesis for the indicated times and lysates were blotted for cIAP2, TRAF1 or as a loading control, β-actin. (**b**) shCtrl or shTRAF1 RAJI cells were treated with 5 µg/ml cycloheximide (CHX) for the indicated times, and lysates were blotted for cIAP2, TRAF1 or as a loading control, β-actin. (**c**) HEK 293T cells overexpressing cIAP2 with or without WT, TRAF1 V203A were treated with cycloheximide (CHX) and blotted, as in panel a. (**d**) HEK 293T cells overexpressing cIAP2 with or without WT, TRAF1 V203A or TRAF1 E207A were treated with cycloheximide and blotted, as in panel a. (**e**) HEK 293T cells overexpressing cIAP2 with or without WT were treated with or without 0.01 mM of the proteasome inhibitor, MG132 or 0.025 mM of the lysosomal inhibitor, chloroquine (CQ). (**f**) HEK 293T were transfected with various combinations of plasmids to overexpress cIAP2, WT TRAF1, cIAP1, cIAP2 H574A, or cIAP1 H588A as indicated and lysates were blotted for cIAP2, TRAF1 or as a loading control, β-actin. All blots are representative of three independent experiments. * non-specific band.

### TRAF1 is needed for cIAP2 recruitment to the TLR4 signaling complex in monocytes

Next, we explored how disrupting the interaction between cIAP2 and TRAF1 would affect inflammatory responses in monocytes. To this end, we employed CRISPR/Cas9 with homologous DNA repair (HDR) to mutate the endogenous TRAF1 in THP-1 monocytes to V203A. Homozygous V203A clones were confirmed with Sanger sequencing (not shown). Crucially, we demonstrated that TRAF1 from V203A mutant THP-1 cells fails to interact with endogenous cIAP2 (**Fig. 3a**), even accounting for the lower cIAP2 levels in the mutant clones. Consistently, cIAP2 protein stability was lower in V203A THP-1 cells compared to WT controls (**Fig. 3b**).

**Figure 3:**
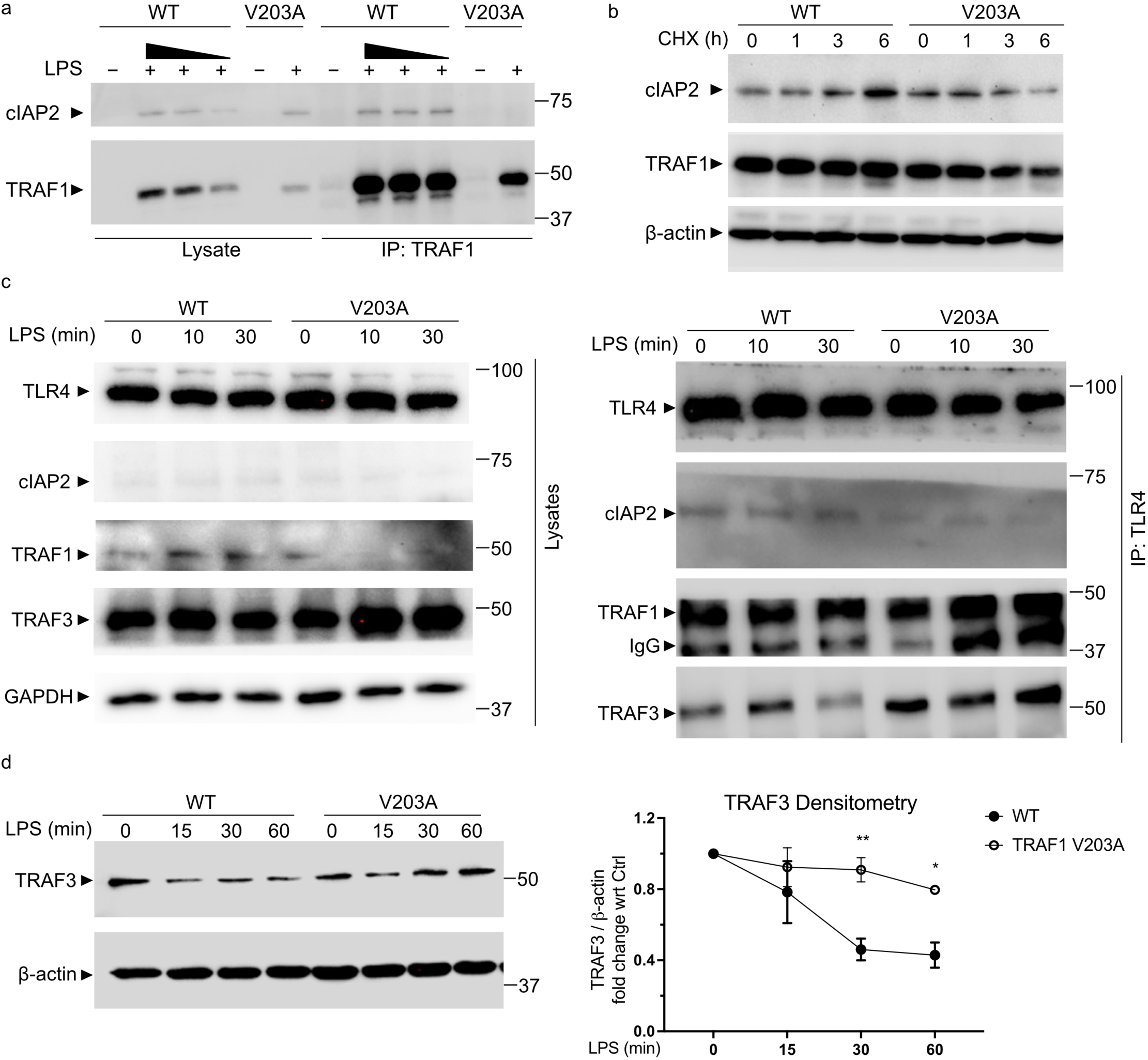
TRAF1 is recruited to TLR4 signaling platform in monocytes. (**a**) WT or V203A TRAF1 mutant THP-1 cells were treated with LPS to induce endogenous TRAF1 and cIAP2 expression followed by Immunoprecipitation (IP) of TRAF1 and immunoblotting for cIAP2 and TRAF1. cIAP2 interaction with WT TRAF1 was maintained despite loading as little as 25% of the lysates from WT (black triangle) compared to V203A. (**b**) WT or TRAF1 V203A THP-1 cells were treated with 5 µg/ml cycloheximide for the indicated times and lysates were blotted for cIAP2, TRAF1 and β-actin. (**c**) PMA differentiated WT or V203A TRAF1 mutant THP-1 cells were stimulated with LPS followed by Immunoprecipitation (IP) of TLR4 and immunoblotting for TLR4, cIAP2, TRAF1, and TRAF3. (**d**) Lysates from LPS stimulated WT or V203A TRAF1 THP-1 cells were blotted for TRAF3 or β-actin as a loading control (left panel). Densitometry of band intensity from 3 independent experiments were quantified by ImageLab software (right panel). All blots are representative of 3 independent experiments. Data in panel d graphs were compared using a two-way ANOVA with multiple comparisons with Sidak correction. *p < 0.05, **p < 0.01.

cIAP2 can be recruited to the TLR4 signaling complex and is required for optimal NF-κB and MAPK (ERK, JNK, Jun) activation. It does so by triggering TRAF3 degradation, which then causes the translocation of the MyD88-associated signaling complex to the cytosol, where TAK1 and its downstream MAPKs are activated. Therefore, we asked whether TRAF1 is important for cIAP2 recruitment to this complex. Indeed, we show for the first time that TRAF1 is part of the TLR4 signaling complex in monocytes, and that TRAF1 V203A mutant cells fail to adequately recruit cIAP2 to the signaling complex (**Fig. 3c**). Importantly, we showed that the reduced levels of cIAP2 in V203A cells reverse the cIAP2-mediated degradation of TRAF3 (**Fig. 3c, d**).

### Endogenous V203A mutation of TRAF1 significantly reduces inflammatory signaling and cytokine production in monocytes

Next, we asked whether the TRAF1/cIAP2 interaction in the TLR4 signaling complex affects LPS-induced inflammation in monocytes. Remarkably, we demonstrated that LPS stimulation of V203A mutant THP-1 cells leads to significantly lower activation of NF-κB than their WT counterparts, as evidenced by reduced degradation of IκBα (**Fig. 4a**), phosphorylation of NF-κB p65 (**Fig. 4a, b**), lower phospho-ERK1/2 signaling (**Fig. 4c**), and attenuated nuclear translocation of NF-κB p65 (**Fig. 4d, e**). Moreover, LPS-stimulated V203A mutant THP-1 cells exhibited diminished activation of TAK1, Jun, and JNK (**Fig. 4f**). Consequently, expression of the pro-inflammatory genes IL-1β, IL-6, and IFN-β were also significantly reduced in V203A cells (**Fig. 4g, Supplementary Fig. 2a**). Similar results were obtained in TRAF1 E207A mutant cells (not shown). Importantly, we show that production of the cytokine IL-2, an indicator of T cell activation, from Jurkat T cells stimulated with anti-CD3 and co-stimulated with anti-4-1BB, is increased when WT TRAF1, but not V203A TRAF1, is overexpressed (**Supplementary Fig. 2b**). Notably, the TRAF1 V203A mutation did not alter the responses of THP-1 cells to TNF stimulation (**Supplementary Fig. 2c-e**).

**Figure 4:**
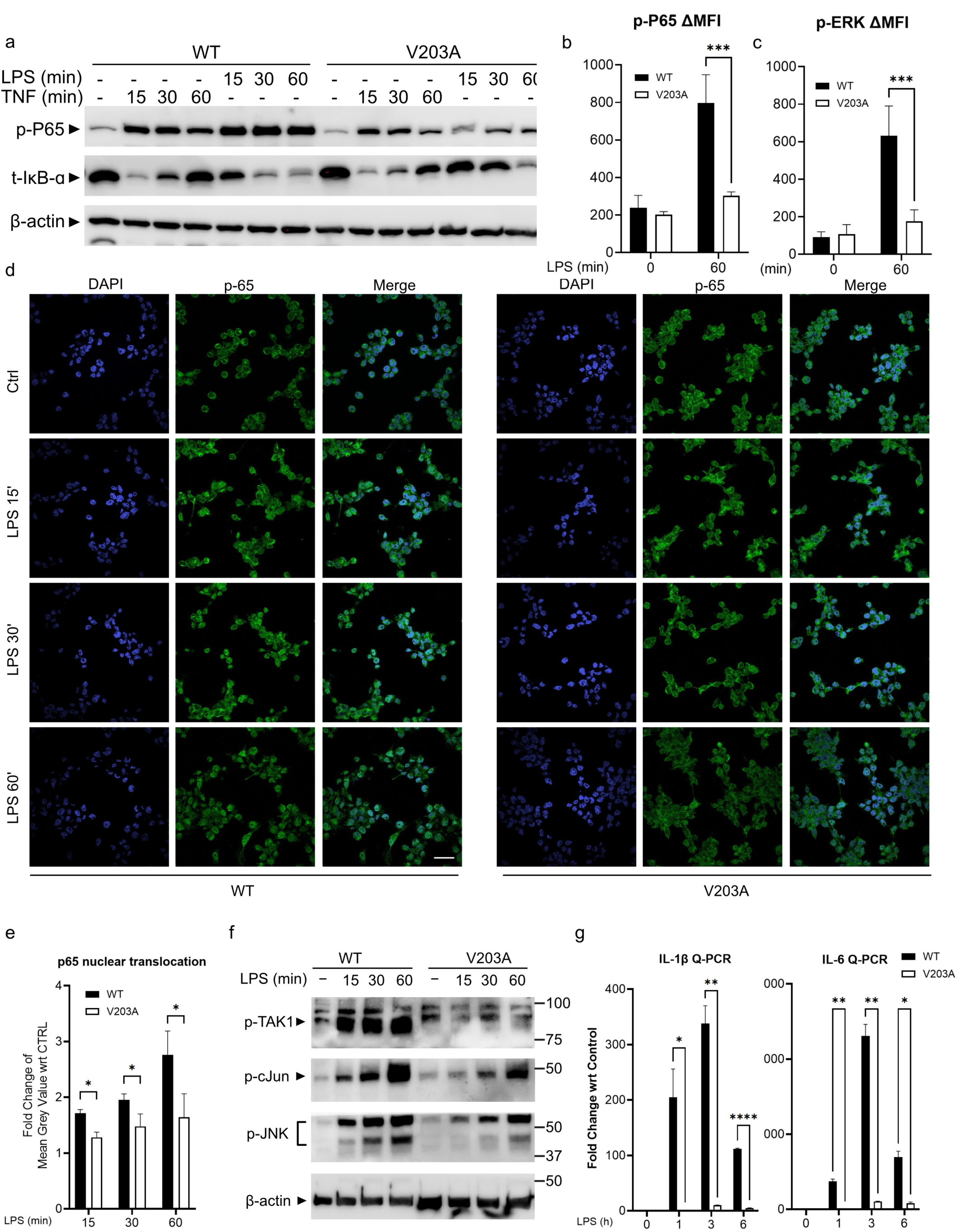
Endogenous V203A mutation of TRAF1 significantly reduces inflammation and cytokine production in monocytes. (**a**) THP-1 cells (WT or V203A) were either stimulated with 100 ng/ml LPS or TNF for the indicated times and lysates were blotted for phospho specific P-65 (p-P65), total IκB-α (t-IκB-α) or β-actin as a loading control. (**b, c**) Mean fluorescence intensities (ΔMFI = MFI of sample − MFI of FMO) of phospho NF-κB p65 (p-P65) (**b**) and phospho ERK1/2 (pERK) (**c**) were measured by flow cytometry in LPS-stimulated WT and V203A THP-1 cells. (**d**) LPS treated WT and V203A TRAF1 THP-1 cells were stained with anti-P65-Alexa Fluor 488 antibody and the nuclear stain, DAPI followed by immunofluorescence (scale bar: 10 μm). (**e**) P65 nuclear translocation was quantified by ImageJ using the overlap of P-65 and DAPI stains from 3 independent experiments as shown in panel c. (**f**) WT and V203A THP-1 cells were stimulated with LPS for the indicated times and lysates were probed for phospho-p-TAK1, p-c-Jun, p-JNK, and as a loading control β-actin. (**g**) Gene expression of IL-1β and IL-6 from LPS-stimulated THP-1 cells was evaluated by real-time Q-PCR. Results were normalized to GAPDH and reported as relative fold change with respect to untreated controls. All blots are representative of 3 independent experiments. Data in all graphs were compared using a two-way ANOVA with multiple comparisons with Sidak correction. *p < 0.05, **p < 0.01, ***p < 0.001, ****p < 0.0001.

### TRAF1 V196A murine macrophages have muted responses to LPS stimulation

We next asked whether disrupting the TRAF1/cIAP2 signaling axis would reduce inflammation in vivo. Human and mouse TRAF1 are highly conserved, and V196 in mice is equivalent to V203 in humans. First, we confirmed that mutating V196 in mouse TRAF1 to alanine (V196A) or glutamate (V196E) abrogated the interaction with cIAP2 (**Fig. 5a**) without affecting the association with HOIL-1 (**Supplementary Fig. 3a**). Next, we generated a CRISPR-mediated knock-in mouse, where the V196A mutation was introduced in TRAF1 (**Supplementary Fig. 3b**). Bone marrow macrophages (BMDMs) from TRAF1 V196A mice failed to recruit cIAP2 to the membrane TLR4 complex, unlike their wildtype (WT) littermate counterparts (**Fig. 5b**). Consequently, LPS stimulation of TRAF1 V196A BMDMs induced significantly lower NF-κB p65 and JNK phosphorylation (**Fig. 5c, d**) and reduced expression of pro-inflammatory cytokines compared to WT littermates (**Fig. 5e, Supplementary Fig. 4a**).

**Figure 5:**
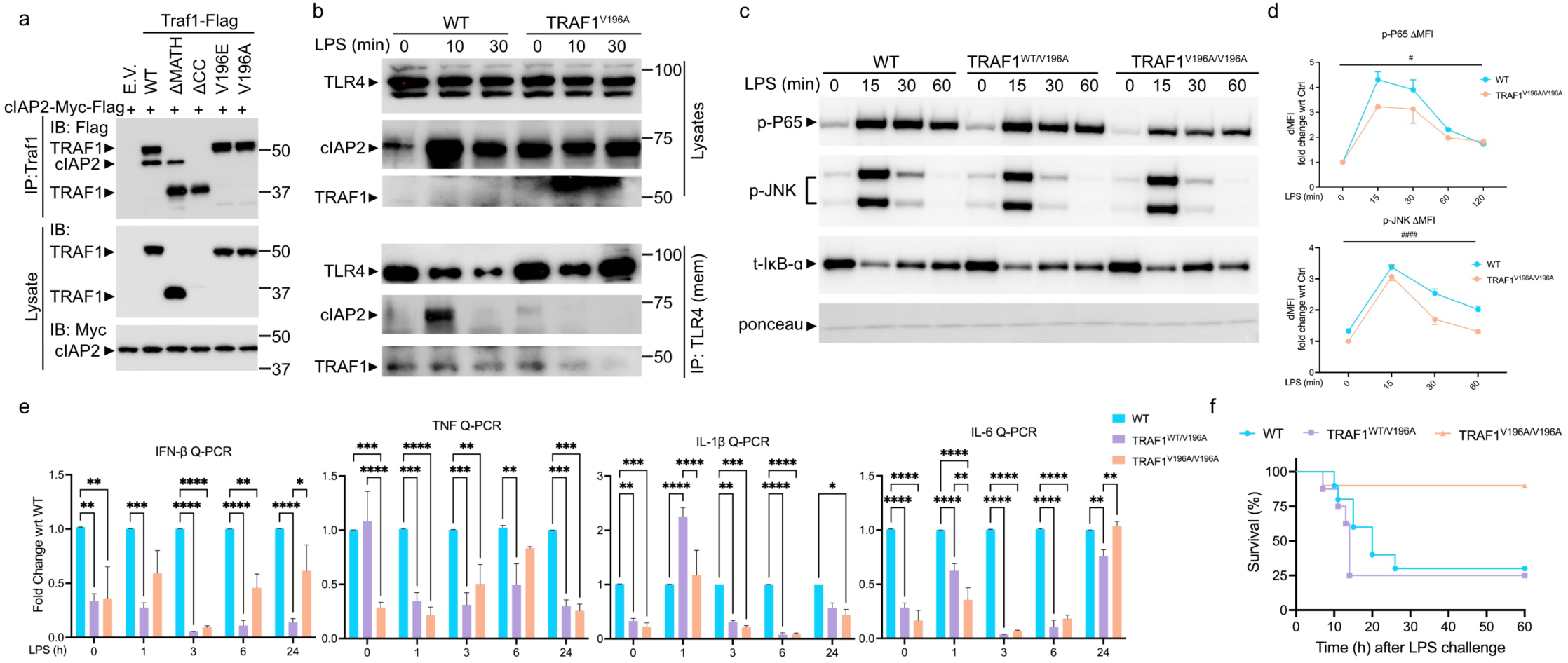
BMDM from TRAF1 V196A mice exhibit reduced responses to LPS and TRAF1 V196A mice are protected from endotoxin-induced septic shock. (**a**) Immunoprecipitation (IP) of mouse TRAF1 from lysates of HEK 293T cells overexpressing mouse cIAP2-Flag-Myc with or without various mouse TRAF1-Flag mutant constructs as indicated and then blotted for Flag, Myc, or TRAF1. (**b**) bone marrow macrophages (BMDMs) were prepared from WT and TRAF1 V196A homozygous mice (TRAF1^V196A^), stimulated with LPS for the indicated times then membrane fractions were prepared (mem) and TLR4 associated proteins were immunoprecipitated then immunoblotted for TRAF1, cIAP2 and TLR4. (c) BMDMs from WT, TRAF1 V196A heterozygous mice (TRAF1^WT/V196A^) and homozygous mice (TRAF1^V196A/V196A^) and stimulated with LPS for the indicated times and immunoblotted for phospho specific P-65 (p-P65), phospho-SAPK/JNK (p-JNK), or total IκB-α (t-IκB-α). Ponceau stain was used to show equal loading. (**d**) Mean fluorescence intensities (ΔMFI = MFI of sample − MFI of FMO) of phospho NF-κB p65 (p-P65) and phospho JNK (pJNK) were measured by flow cytometry. (**e**) Gene expression of IFN-β, TNF, IL-1β, and IL-6 from LPS-stimulated WT, TRAF1 V196A heterozygous and homozygous BMDMs was evaluated by real-time Q-PCR. Results were normalized to RPLP0 and reported as relative fold change with respect to WT controls. Data are representative of at least 3 independent experiments. Data in all graphs were compared using a two-way ANOVA with multiple comparisons with Sidak correction. *p < 0.05, **p < 0.01, ***p < 0.001, ****p < 0.0001. ^#^p<0.05, ^####^p<0.0001 for Anova of difference between WT and TRAF1^V196A/V196A^ groups. (**f**) Survival of wildtype (WT), heterzygous (TRAF1 V196A/WT) and (TRAF1 V196A/V196A) littermates challenged with LPS (35 mg per kg body weight intraperitoneally), monitored over time. n = 10 mice per group from two independent experiments of male and female mice.

### TRAF1 V196A mice are protected from sepsis

Macrophages are early responders to microbial presence, using TLRs to sense pathogens and promote inflammatory responses. In some cases, especially if the pathogen becomes bloodborne, TLR4 stimulation by LPS causes a systemic and exuberant response leading to a cytokine storm and sepsis. Therefore, we asked whether the V196A TRAF1 mutation alters LPS-induced septic shock in mice. Remarkably, while most WT littermate mice succumbed to LPS-induced sepsis, most of the homozygous TRAF1 V196A mice survived (**Fig. 5f**). V196A heterozygous mice (TRAF1 WT/V196A) were not protected (**Fig. 5f**), indicating that a single copy of TRAF1 is sufficient to sustain TLR4 signaling. Surprisingly, there were no significant differences in pro-inflammatory cytokine production in the serum of V196A mice compared to WT littermates following a sublethal dose of LPS (**Supplementary Fig. 4b**). These results demonstrate that disruption of the TRAF1/cIAP2 axis protects mice from LPS-induced sepsis.

### TRAF1 V196A mice exhibit reduced joint inflammation in a model of RA

After showing that blocking the association of TRAF1 with cIAP2 lowers macrophage inflammation and protects against sepsis, we asked whether targeting this signaling axis could be beneficial in a complex autoimmune disease like rheumatoid arthritis. To this end, we employed the well-established collagen antibody-induced arthritis (CAIA) model in WT and TRAF1 V196A littermates. Most studies using this model administer LPS on day 3 to trigger an aggressive disease course (*25*, *26*). However, given that we showed V196A mice have attenuated responses to LPS, we modified the model to only use the collagen antibody cocktail (two intraperitoneal injections of ArthritoMab cocktail on days 0 and 1), which triggered moderate levels of systemic joint inflammation (*23*). Remarkably, V196A mice exhibited significantly reduced knee and ankle joint inflammation compared to WT littermates (**Fig. 6a, b**). Moreover, rear paw thickness and mean arthritis scores were also markedly lower in TRAF1 V196A mice (**Fig. 6c, d**). On day 14, inflammatory cell infiltrates and bone erosion in the knee joints of TRAF1 V196A mice were significantly lower than those in WT littermates (**Fig. 6e**).

**Figure 6:**
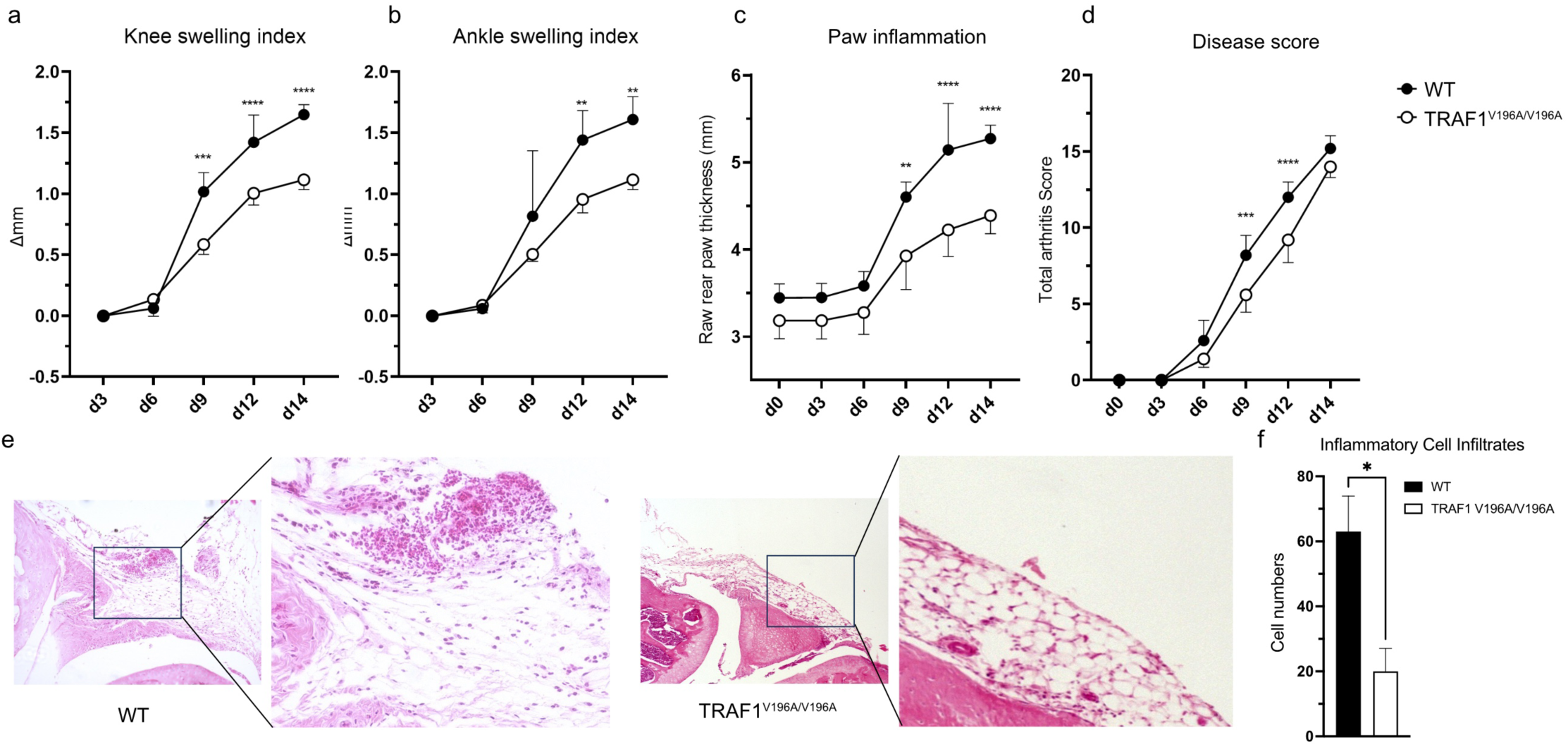
TRAF1 V196A mice show reduced joint inflammation and cellular infiltration following collagen antibody induced arthritis (CAIA) model. WT and TRAF1^V196A/V196A^ littermate mice were injected intraperitoneally with 5 mg ArthritoMab^TM^ Antibody Cocktail (MD Bioproducts) on day 0 and 5 mg on day 1. Knee (**a**) and ankle (**b**) joint thickness, and paw thickness (**c**) were measured on day 0, then on days 3, 6, 9, 12 and 14 using a digital caliper. Data was reported as the change in thickness (1′ mm). (**d**) Total arthritis score from mice in panels a-c was reported by scoring all four paws were scored on a scale of 0-4 and combining the scores each day. (**e**) Histopathological evaluation of mice showing representative H&E staining images for synovial tissue of the knee joint sections at d14. (**f**) Quantification of the average inflammatory infiltrates per section from panel e. n=5 per genotype. Statistical analysis was performed using two-way ANOVA with multiple comparisons with Sidak correction. *p < 0.05, **p < 0.01, ***p < 0.001, ****p < 0.0001.

Overall, we show for the first time that selectively disrupting TRAF1’s interaction with cIAP2 attenuates inflammatory signaling in monocytes and macrophages, leading to protection from sepsis and reduced severity of RA. These findings may ultimately pave the way for novel therapeutics for RA.

## DISCUSSION

RA, though multifactorial, is characterized by misdirected and overactive immune/inflammatory responses leading to joint damage. The mechanisms driving RA pathogenesis are not completely understood. TRAF1, a key adapter protein for immune cell signaling, plays a paradoxical, context-dependent role in both blocking NF-κB signaling to limit inflammation in innate immune cells (e.g., macrophages) (*15*, *24*) and promoting activation, proliferation, and cytokine production in adaptive lymphocytes (T and B cells) (*8*, *14*, *27*). The dual role of TRAF1 depends on its ability to recruit cIAP2 to signaling complexes, promoting NF-κB and MAPK activation (*14*, *27*), while also directly interfering with the linear ubiquitin chain assembly complex (LUBAC) to limit NF-κB activation downstream of TLR signaling (*15*). Therefore, studying the role of TRAF1 in complex diseases that rely on various signaling pathways and different immune cell types is challenging. We hypothesized that selectively targeting TRAF1/cIAP2 signaling would likely attenuate inflammatory diseases. TRAF1 polymorphisms have been associated with increased risk of RA (*16*, *17*, *21*), and recent studies have shown that lower TRAF1 levels promote inflammation and worsen joint damage in gout (*24*). Moreover, injection of TRAF1-deficient macrophages into the knee joints of mice subjected to the CAIA model of RA exacerbated inflammation (*23*). To break TRAF1’s dichotomous role in NF-κB signaling, we pinpointed the site of interaction between TRAF1 and cIAP2 to valine 203 (valine 196 in mice) and showed that TRAF1 V203A mutant THP-1 cells exhibit markedly attenuated inflammatory responses to LPS but not TNF stimulation. We showed that TRAF1 is recruited to the TLR4 complex, where it protects cIAP2 from degradation by cIAP1, which in turn causes the degradation of TRAF3 and subsequently promotes the activation of TAK1 and downstream NF-κB and MAPK (ERK, JNK) signaling. Remarkably, V196A mice, which similarly fail to recruit cIAP2 to the membrane TLR4 signaling complex, exhibited markedly reduced inflammatory signaling in response to LPS in vitro and were protected from LPS-induced sepsis in vivo. Interestingly, despite this protection, we observed no significant changes in cytokine secretion into the serum in the sublethal model of sepsis. This unexpected finding could be attributed to the dose and timing of cytokine measurements, which may not have captured transient changes in cytokine levels. Finally, in a model of RA, V196A mice showed significantly lower inflammatory cell infiltration and swelling in the joints and paws compared to their WT littermates.

Current therapies for RA rely on biologics to block the action of specific cytokines or general immunosuppressive drugs. However, these strategies are ineffective in about half of the patients, who often develop resistance after being switched to second-line biologics (*28*). Therefore, there is a need for a new class of inhibitors that can suppress the production of several cytokines that promote RA without dampening the entire immune response. TRAF1 represents an excellent opportunity for such an approach, as we have shown that selectively targeting its interaction with cIAP2 attenuates inflammatory responses and reduces the severity of inflammation-driven diseases such as sepsis and RA. Designing new therapies that specifically interfere with the cIAP2-TRAF1 interaction could lead to a new generation of drugs for RA therapy.

## Author contributions

AAS conceived, designed, and interpreted experiments and wrote the manuscript, with input from YT, FA and SSM. YT, FA, SSM and AM conducted experiments with the help of SZ, JT and SS. RI and GS provided lab resources and equipment to conduct some experiments.

## Materials and Methods

### Cell Culture and Reagents

Day 5 primary bone marrow derived macrophages (BMMs) were prepared from C57BL/6 and TRAF1 V196A mice as previously reported (*15*, *24*). THP-1, RAJI, and 293T/17 cells were obtained from the American Type Culture Collection (ATCC). All cells were cultured in RPMI 1640 (Sigma Aldrich) supplemented with 10% FCS, 2-ME, sodium pyruvate and non-essential amino acids (Gibco). LPS (*Escherichia coli*; serotype O26:B6) and cycloheximide were from Sigma Aldrich. Recombinant human TNF was from Invitrogen.

### Crispr/Cas9 editing of TRAF1

THP-1 cells were electroporated (Nucleofactor 4D; Lonza) with an RNP complex comprised of AltR Cas9 enzyme (Integrated DNA technologies; IDT) and a crRNA: tracer RNA duplex (sgRNA) targeting TRAF1 near V203 (crRNA: AAGCTGCGTGTGTTTGAGAACATTGTTGCTGCCCTCAACAAGGAGGTGGAGGCCTCC). Cells were then cultured in media containing Alt-R HDR enhancer (IDT) and cleavage was confirmed by a T7-cleavage assay (IDT). Cells were then cloned by limiting dilution and individual clones were screened for V203A mutation by the Guide-it Knock-in Screening Kit (Takara Bio). Homozygous clones were sequenced by Sanger sequencing to confirm the presence of a homozygous mutation without any off-target effects.

### Subcellular fractionation and co-immunoprecipitation

lysates were prepared from 2 × 10^6^ THP-1 or immortalized bone marrow derived macrophages (iBMDM) cells. Membrane and cytosolic fractions were separate by Mem-PER™ Plus Membrane Protein Extraction Kit (Thermo Fisher) following manufacturer’s instruction and immunoprecipitated with 1.5 μg of anti-TRAF1 antibody (clone 1F3; erviceeinheit Monoklonale Antikörper; Institut für Molekulare Immunologie, Munich, Germany) and then resolved on 10% SDS gel before being probed with indicated antibodies.

### Immunofluorescence

THP-1 cells were plated in 24 well-plates containing glass slides and differentiated by PMA (100 nM; overnight) and then stimulated with LPS followed by staining by DAPI (Thermo-Fisher) and anti-p65 antibody (Cell Signaling), mounted on slides, imaged by immunofluorescence on a confocal microscope (Eclipse Ti2 confocal microscopy, Nikon) and quantified by Image J software (National Institutes of Health).

### RNA isolation and Real-time PCR

mRNA was isolated from the THP-1 cells using Trizol reagent (Thermo Fisher) following manufacturer’s instruction. Total RNA was converted to cDNA by standard reverse transcription with M-MuLV Reverse Transcriptase (New England Biolabs, Ipswich, MA) and oligo(dT) primers (Qiagen, Hilden, Germany), and then amplified with a Ssoadvanced SYBR Green master mix (Bio-Rad) and human primers (listed below) using a CFX384 Touch Real-Time PCR Detection System (Bio-Rad). The primers used are listed in *Supplementary Table 1*. Real time PCR included initial denaturation at 95°C for 10 min, followed by 40 cycles of 95°C for 30 s, 55°C for 1 min, 72°C for 1 min, and one cycle of 95°C for 1 min, 55°C for 30 s, 95°C for 30 s. Fold change in gene expression was calculated using the delta-delta Ct (2^−ΔΔCt^) method with respect to untreated (PBS; cp) controls. Data are averages from three biological replicates and variations reported as standard deviations.

### Mice and models of sepsis and RA

V196A mice were generated in the C57BL/6 background at the Toronto Centre for Phenogenomics (TCP) where Cas12a(Cpf1) was employed to target TRAF1 to cause TG>CA mutation using the gRNA sequence: CAAACATTGTTGCTGTCCTCA. Mice were validated for absence of off-target effects and TRAF1 was sequenced to ensure the presence of the desired mutation. C57BL/6 Wild type mice (Charles River) were crossed with TRAF1 V196A mice and F1 progeny were bred for all in vivo experiments. Mice used for the in vivo LPS challenge were littermates between 6–8 weeks old. LPS from *Escherichia coli* (serotype O26:B6; Sigma Aldrich) was diluted in sterile PBS and injected intraperitoneally (200 μl) into mice as indicated in figure legends. Mice used for the collagen induced arthritis model were injected with 5 mg of ArthritoMab^TM^ Antibody Cocktail (MD Bioproducts) on day 0 and another 5 mg (i.p.) on day 1 (*23*). We opted against using LPS to study TLR4 independent effect of TRAF1. Mice were then monitored daily to score for mean arthritis, and knee, ankle and paw thickness was measured by a caliper on days 0, 3, 6, 9, 12 and 14. All animals were housed under specific pathogen free conditions at the Farquharson building vivarium at York University and all procedures were approved by York University Animal Care Committee in accordance with Canadian Council on Animal Care regulations (protocol 2019-12 for CAIA and 2023-06 for LPS sepsis).

### Statistical analyses

All statistical analysis was done using GraphPad software (Prism) using 2 way analysis of variance (ANOVA) for comparison of multiple groups, with p-values as indicated in the figure legends. For the murine sepsis model, differences in survival were analyzed by the Log-rank (Mantel-Cox) Test.

## Acknowledgements

Grant support: This work was supported by the Canadian Institutes of Health Research (201809PJT), the Arthritis Society (18–0276), and a discovery award from the Banting Foundation.

---

**Supplemental Table 1.**
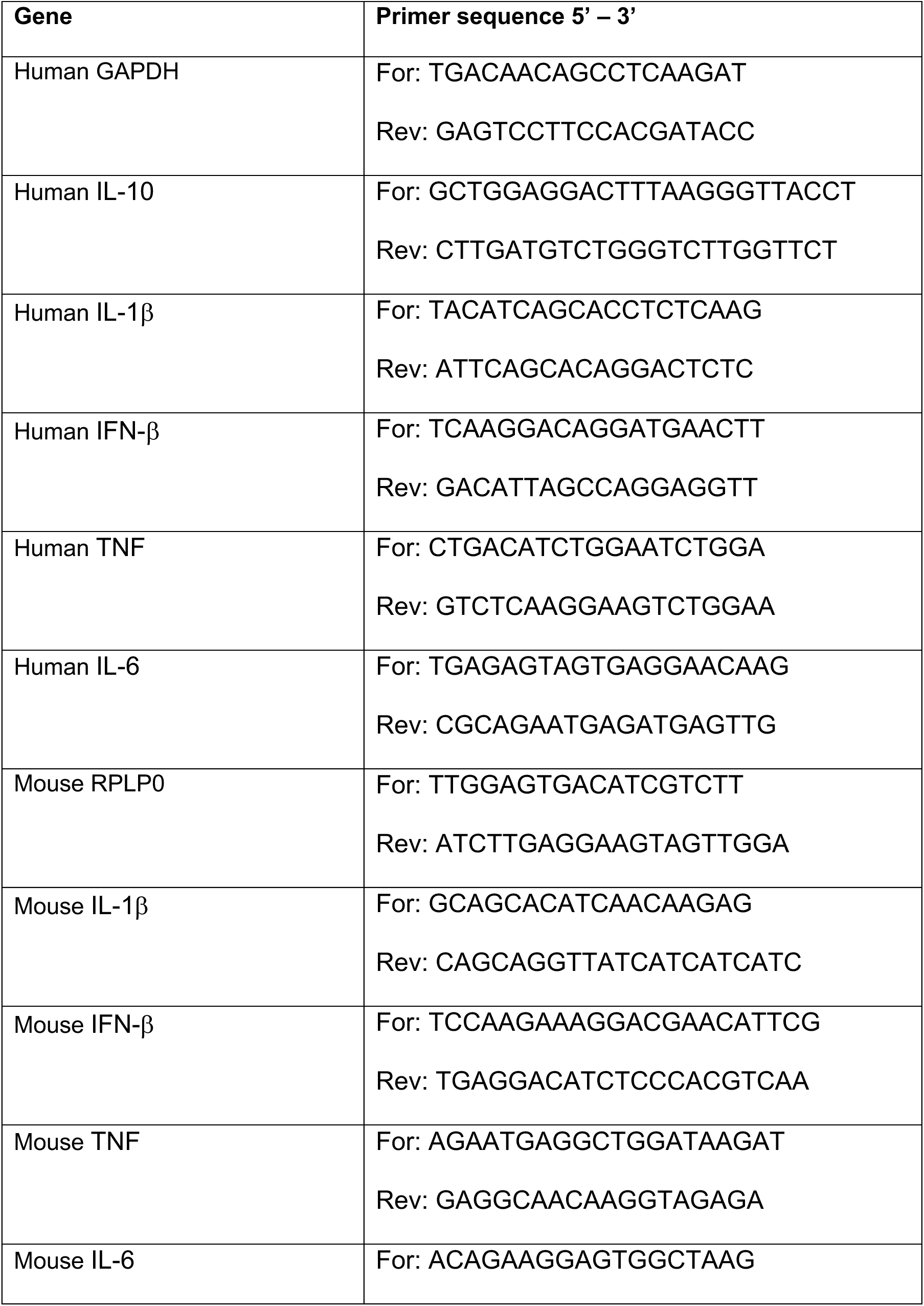
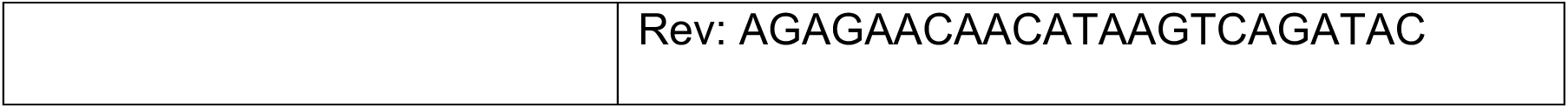

## Supplemental Figure Legends

**Supplemental Figure 1:**
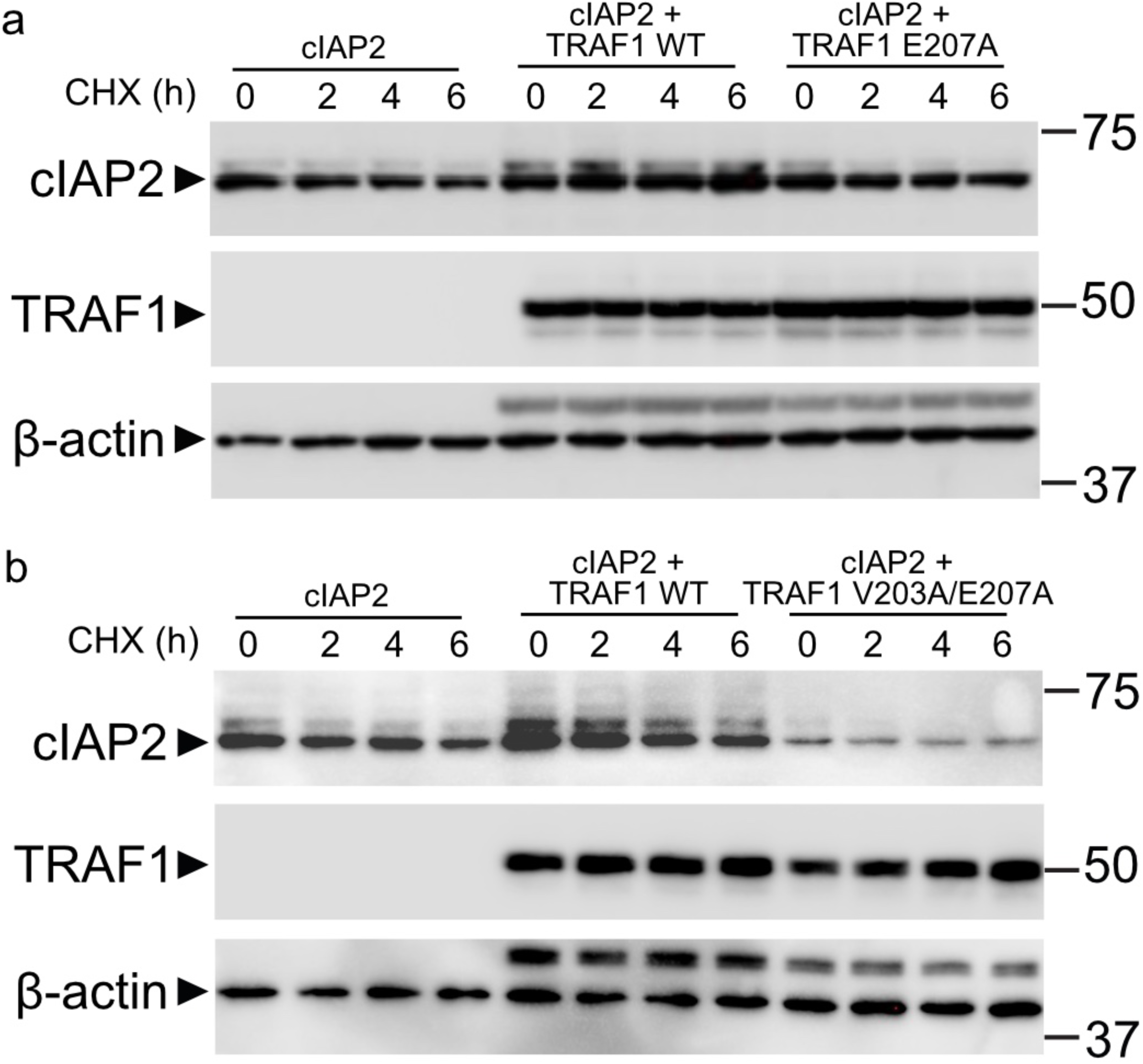
(**a**) Immunoprecipitation (IP) of TRAF1 from lysates of HEK 293T cells overexpressing cIAP2 with or without wildtype (WT) or E207ATRAF1 mutant constructs as indicated and then treated with cycloheximide and lysates were blotted for cIAP2, TRAF1, or as a loading control β-Actin. (**b**)) Immunoprecipitation (IP) of TRAF1 from lysates of HEK 293T cells overexpressing cIAP2 with or without wildtype (WT) or V203A/E207A TRAF1 double mutant constructs as indicated and then treated with cycloheximide and lysates were blotted for cIAP2, TRAF1, or as a loading control β-Actin.

**Supplemental Figure 2:**
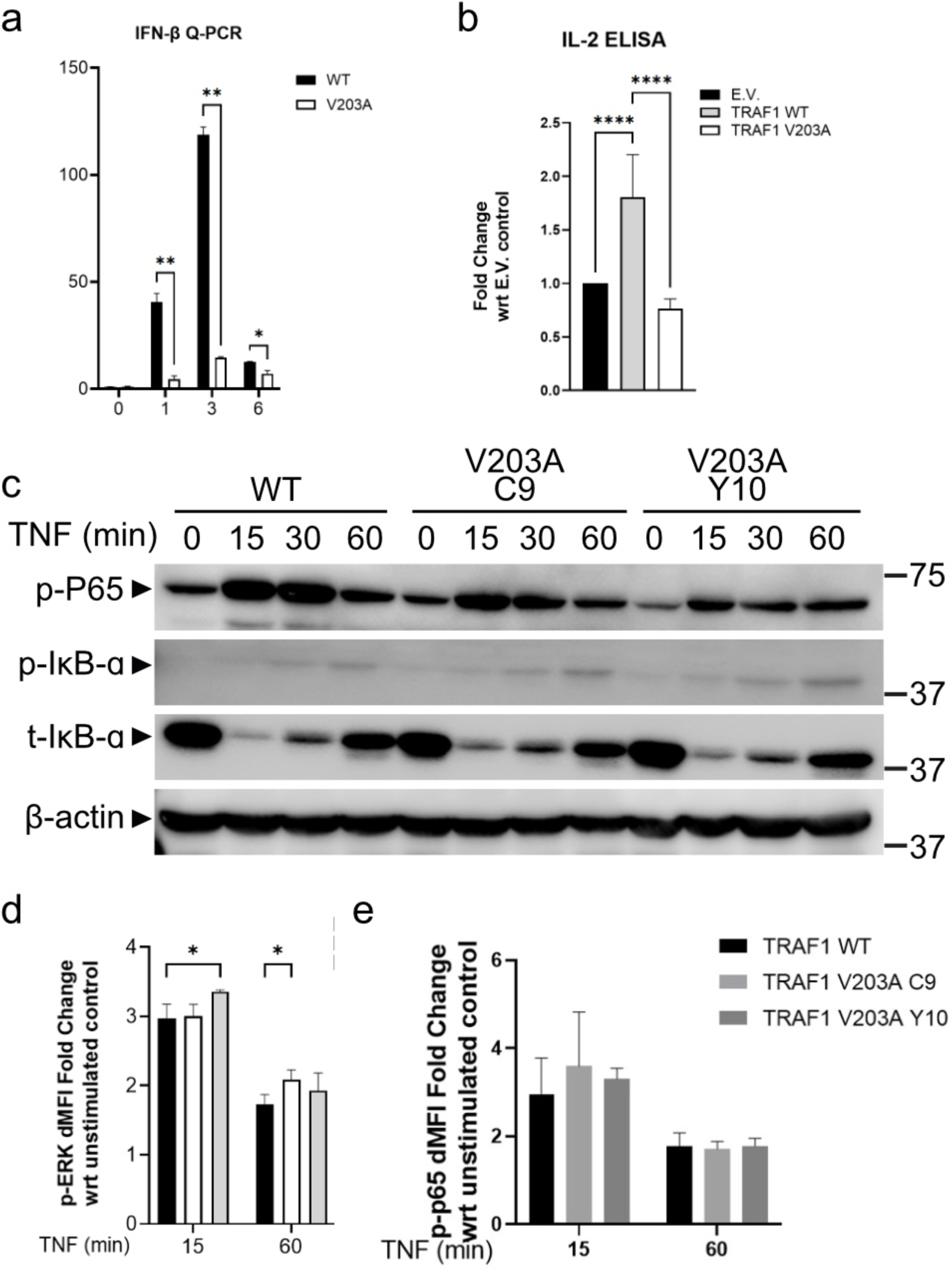
(**a**) Gene expression of IFN-β from LPS-stimulated WTY or V203A THP-1 cells was evaluated by real-time Q-PCR. Results were normalized to GAPDH and reported as relative fold change with respect to untreated controls. (**b**) Jurkat cells stably expressing 4-1BB were transfected with an empty vector (E.V.), TRAF1 WT or TRAF1 V203 plasmids and then stimulated with anti-CD3 + anti-41BB agonist Mabs for 24 h and IL-2 from supernatants was measured by ELISA. (**c**) THP-1 cells (WT or V203A) were stimulated with 100 ng/ml TNF for the indicated times and lysates were blotted for phospho specific P-65 (p-P65), phospho IκB-α (p-IκB-α), total IκB-α (t-IκB-α) or β-actin as a loading control. (**d, e**) Mean fluorescence intensities (ΔMFI = MFI of sample − MFI of FMO) of (**d**) phospho ERK1/2 (pERK) and (**c**) phospho NF-κB p65 (p-P65) were measured by flow cytometry in TNF-stimulated WT and V203A THP-1 cells.

**Supplemental Figure 3:**
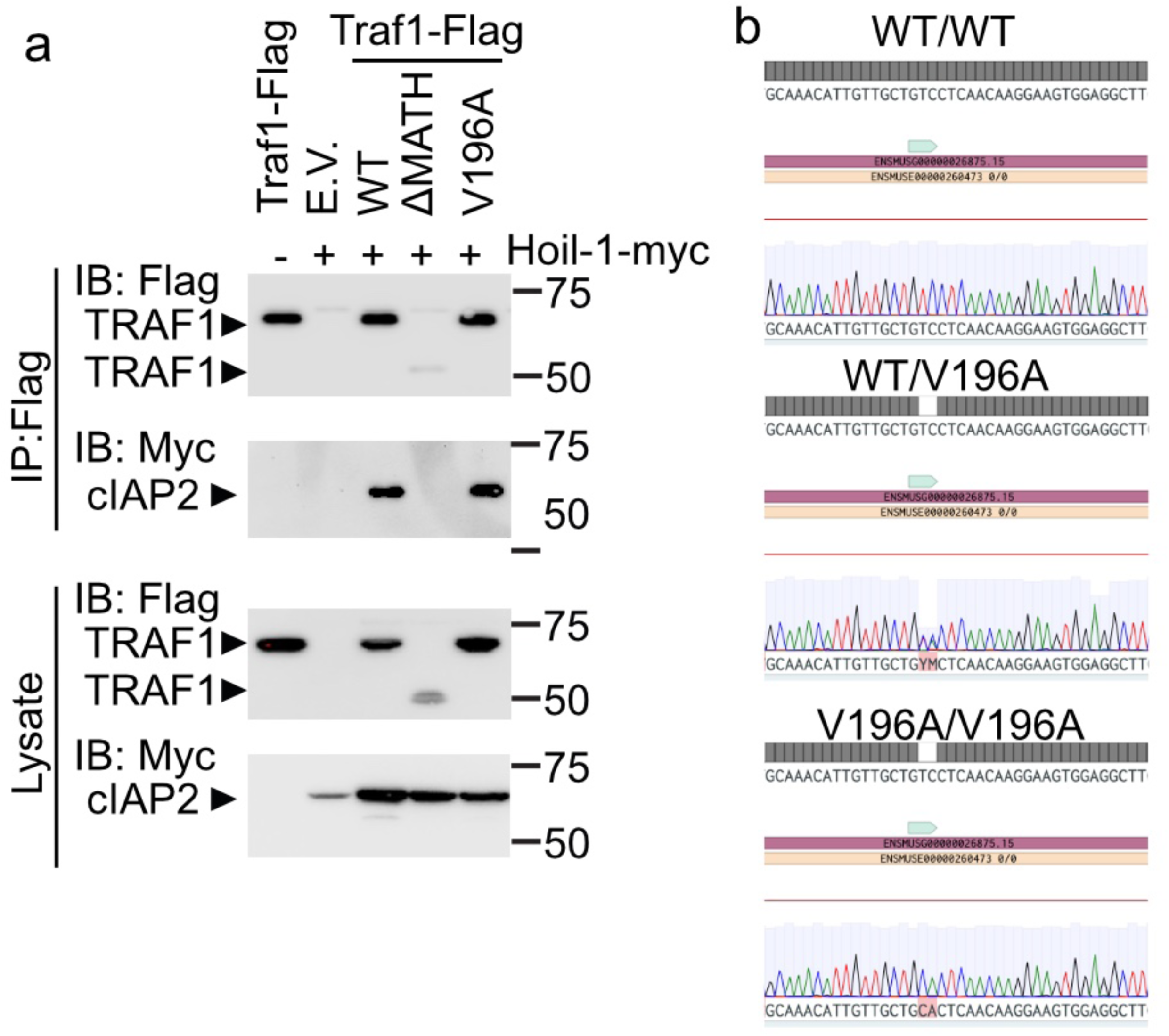
(**a**) Co-immunoprecipitation of mTRAF1 lysates of HEK293T/17 cells, followed by immunoblotting (IB) with Myc and Flag specific antibodies and light-chain-specific anti-Rabbit- and light-chain-specific anti-mouse. This experiment was conducted three independent times. (**b**) Sanger DNA sequencing of the region flanking the site of mutation in TRAF1 gene from WT, heterozygous (TRAF1 WT/V196A) and homozygous (TRAF1 V196A/V196A) knockin mice.

**Supplemental Figure 4:**
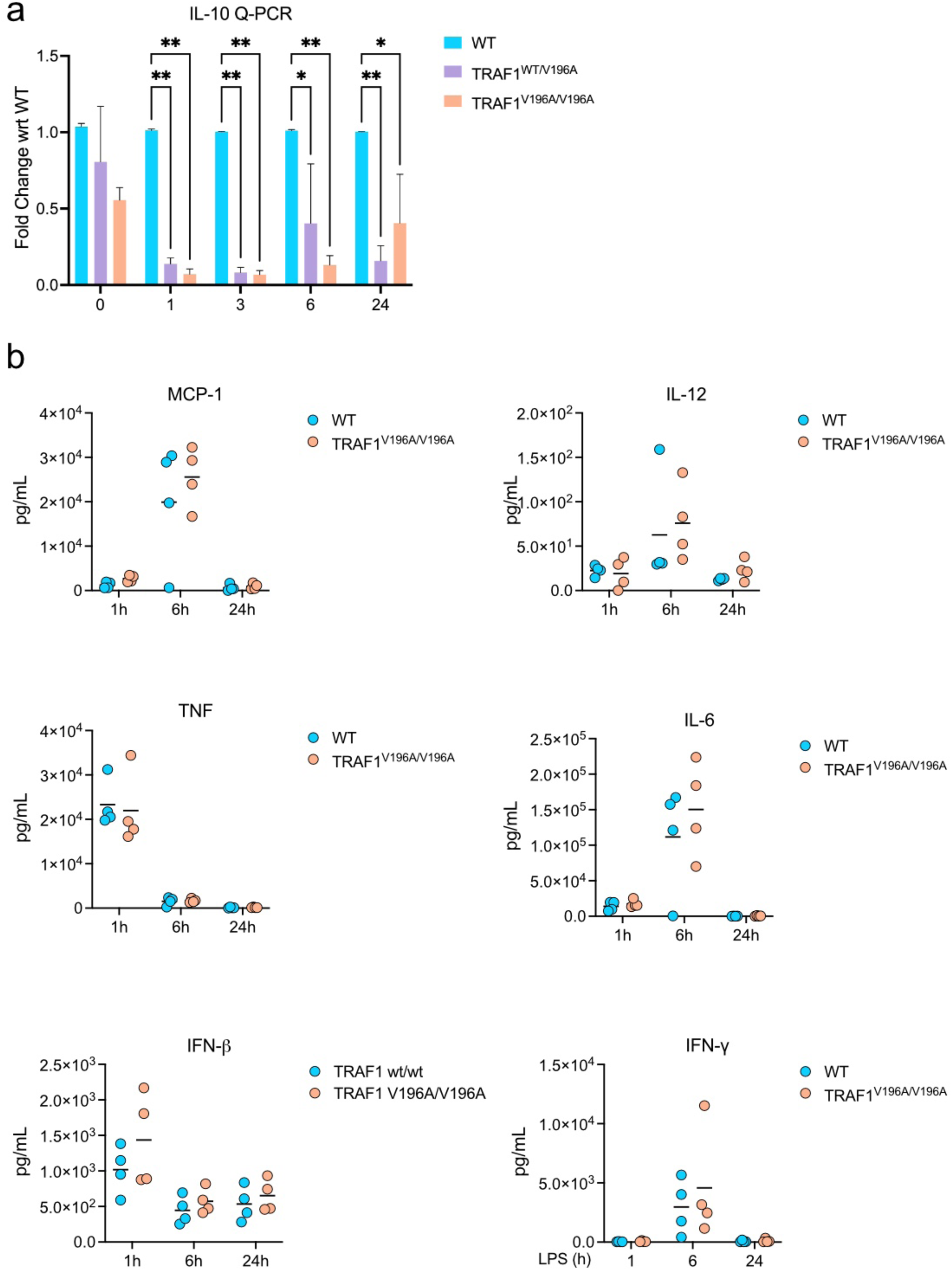
(**a**) Gene expression of IL-10 from LPS-stimulated WT, TRAF1 V196A heterozygous and homozygous BMDMs was evaluated by real-time Q-PCR. Results were normalized to RPLP0 and reported as relative fold change with respect to untreated controls. (**b**) Multiplex cytokine assay of TNF, MCP-1, IL-12 p70, IL-6, IFN-γ, and IFN-β in serum from WT, V196A/WT and V196A/V196A littermate mice, assessed 1, 6 and 24 h after intraperitoneal injection of a sublethal dose of LPS (10 mg per kg body weight). Each symbol represents one mouse; n = 4 mice of each genotype. Statistical analysis was performed using two-way ANOVA with multiple comparisons with Sidak correction. *p < 0.05, **p < 0.01.

